# Variation in white spruce needle respiration at the species range limits: a potential impediment to northern expansion

**DOI:** 10.1101/2021.08.18.456715

**Authors:** Kevin L. Griffin, Zoe M. Griffin, Stephanie C. Schmiege, Sarah G. Bruner, Natalie T. Boelman, Lee A. Vierling, Jan U. H. Eitel

**Author notes:** Correspondence: Kevin L. Griffin.

## Abstract

White spruce (*Picea glauca*) spans a massive range from arctic treeline to temperate forests, yet the variability in respiratory physiology and related implications for tree carbon balance at the extremes of this distribution remain as enigmas. Working at both the most northern and southern extents of the white spruce distribution range more than 5000 km apart, we measured the short-term temperature response of dark respiration (*R*/T) at upper and lower canopy positions. *R*/T curves were fit to both polynomial and thermodynamic models so that model parameters could be compared among locations, canopy positions, and with previously published data. Respiration measured at 25°C (*R*25) was 68% lower at the southern location than at the northern location (0.73±0.15 vs. 2.27±0.02 μmol m^-2^ s^-1^), resulting in a significantly lower (p< 0.01) intercept in *R*/T response in temperate trees. Only at the southern location did upper canopy leaves have a steeper temperature response than lower canopy leaves, likely reflecting steeper canopy gradients in light. No differences were observed in the maximum temperature of respiration. At the northern range limit respiration is nearly twice that of the average *R*25 reported in a global leaf respiration database. This large carbon cost likely challenges tree survival and contributes to restricting the location of the northern treeline. We predict that without significant thermal acclimation, foliage respiration will increase with projected end-of-the-century warming and will likely constrain the future range limits of this important boreal species.

**Summary Statement:** White spruce (*Picea glauca*) needle respiration at the northern limit of the species range is three times higher than at the southern range limit (when measured at 25 °C). This high carbon cost likely challenges tree survival and contributes to the location of the northern treeline.

## Introduction

The distribution range of a species delineates the geographical location where historical, physiological and biotic filters combine to result in successful growth and reproduction (Lambers & Oliveira, 2019). The niche-breadth hypothesis explains species distributions based on envelopes of environmental conditions tolerated (Lowry & Lester, 2006), yet there is no universally accepted “cause” for delineating a species’ range despite a long history of debate (see Casazza et al., 2005; Kruckeberg & Rabinowitz, 1985; Kunin & Gaston, 1993; Lavergne et al., 2004; Stebbins, 1942; Watson, 1833). Clearly many other evolutionary and ecological factors influence species range distributions, including genetic diversity, phenotypic plasticity, reproduction strategies, and metapopulation dynamics (*reviewed in* Brown et al., 1996; Gaston, 1996; Lowry & Lester, 2006). Still, the role of the growth environment is undeniable and links a species’ distribution to its physiological performance. Using a physiological approach to understand the complex relationships between climate and the distribution of species can be preferable to simple climate envelope models because the former is capable of predicting species distributions under a variety of possible environmental conditions (Hijmans & Graham, 2006; Malanson et al., 1992; Prentice et al., 1992), including novel conditions that are likely to occur with rapid climate change.

Physiological processes such as photosynthesis, respiration and growth all respond strongly to local environmental conditions. Together these processes constrain plant carbon balance and thus contribute to a species distribution range (part of the physiological filter of Lambers & Oliveira, 2019). Here, we concentrate on leaf respiration. This flux is less well studied than photosynthesis, provides a crucial link between photosynthesis (carbon gain) and growth (carbon sequestration) and has been hypothesized to control the range distribution of individual species (Criddle et al., 2003). Furthermore, respiration is highly temperature sensitive, making it an important determinant of ecosystem productivity (Valentini et al., 2000). *GlobResp* (Atkin et al., 2015), a global database of plant respiratory characteristics, identifies latitudinal gradients in leaf respiration measured at a common temperature and these respiration gradients increase with absolute latitude. These findings suggest that species with a large range should exhibit variable rates of respiration across their distribution, although this hypothesis has not been explicitly tested (Atkin et al., 2015). Patterson et al. (2018) quantified respiratory rates and responses to temperature in 16 tree species growing in a common temperate location, grouping them by their relative location within their individual species distributions. The results show that northern ranged species growing near their southern range limit had 71% higher respiration rates (measured at 20 °C) than southern ranged species growing near the northern edge of their range limits. Quantifying respiratory characteristics of individuals growing near the margins of their species range distributions can elucidate physiological controls of the current distribution.

Recently, we reported that white spruce (*Picea glauca* (Moench) Voss) growing at the arctic treeline (which marks the transition between the boreal forest and treeless tundra occurring in the Forest Tundra Ecotone (FTE)) exhibit high respiratory costs (Griffin et al., 2021). As one of the hardiest coniferous species, white spruce has a suite of structural and functional traits that are adapted to cold temperatures and short growing seasons (Sutton, 1969) . With a transcontinental range in North America from the west coast of Alaska to the east coast of Canada and New England (US Geological Survey, 1999) (Figure 1), white spruce is one of the most common tree species defining the FTE. Perhaps surprising given this large geographical range, several studies show that white spruce has limited physiological plasticity (Benomar et al., 2018; Man & Lieffers, 1997; McNown & Sullivan, 2013; Prud’Homme et al., 2018; Stinziano & Way, 2017; Weger & Guy, 1991). However, McNown & Sullivan (2013), working across a gradient that included terrace, forest and treeline sites, demonstrated that the physiological capacity of white spruce can vary due to other site factors such as soil properties and nutrition, even when climatic variables like temperature are constant. Furthermore, ambient environmental conditions at the southern edge of its range distribution can be markedly different from those at northern edge, implying that white spruce can be successful under vastly different environmental extremes. To our knowledge no studies have compared the respiratory characteristics of this species at the opposite ends of its distribution where climate and light conditions differ dramatically.

**Figure 1.**
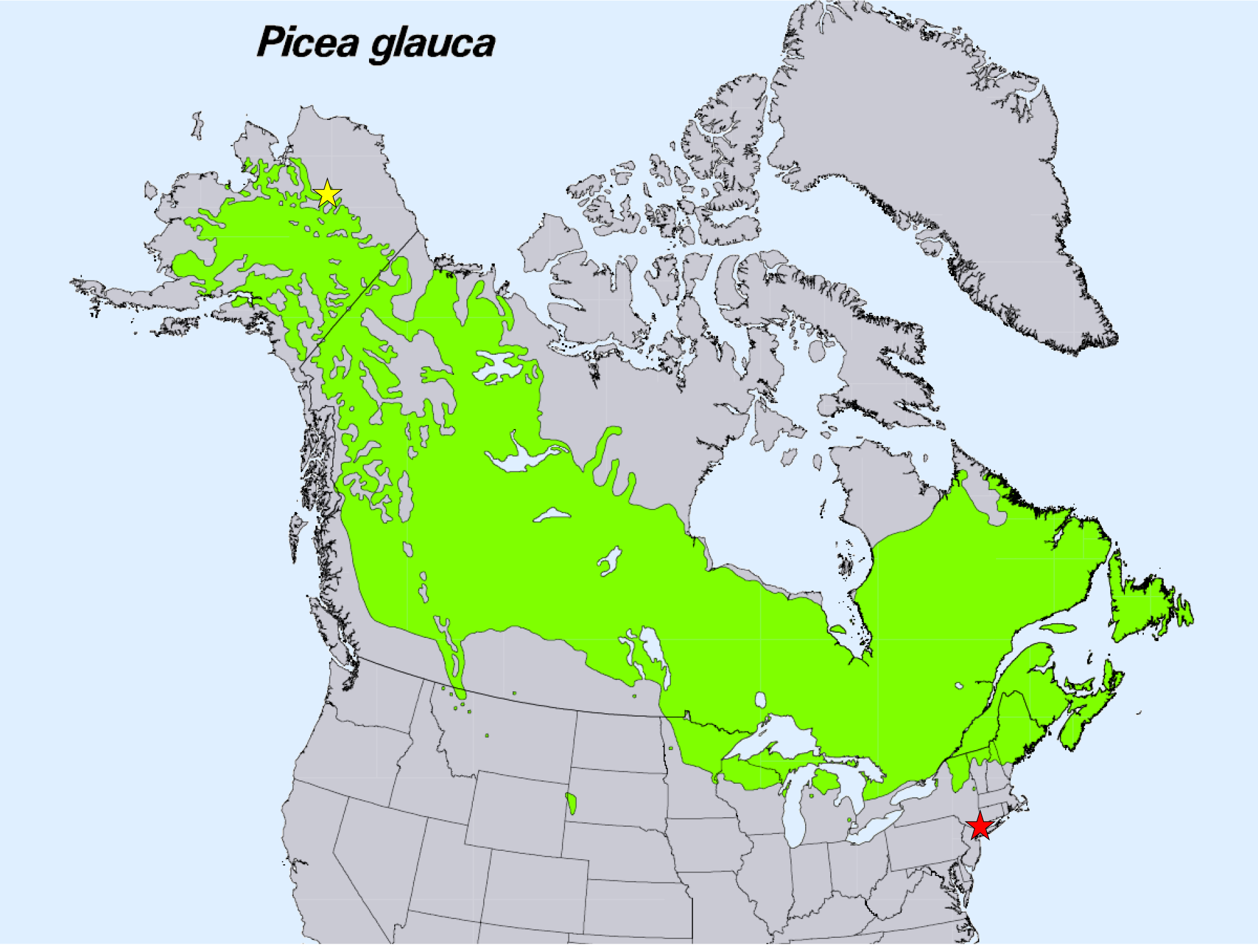
Species range distribution for *Picea glauca.* The southern range limit site, Black Rock Forest, Cornwall NY, is marked with the red star. The arctic treeline site in northern Alaska is marked with a yellow star. Map from U.S. Geological Survey, Department of the Interior/USGS based on original data from Little (2004).

Canopy position can also significantly influence leaf respiration (Griffin et al., 2001, 2002; Tissue et al., 2002; Whitehead et al., 2004; Xu and Griffin 2006; Weerasinghe et al., 2014; Araki et al., 2017). Generally, the top of the canopy is the most physiologically active portion of a tree crown and this has been shown to affect both average respiration rates and the response of respiration to temperature (Griffin et al., 2001, 2002; Araki et al., 2017). In addition, substantial environmental gradients can exist from the top to the bottom of a tree canopy and these should also have predictable effects on foliar respiration. The strength of these canopy environmental gradients is affected by physical factors influencing the available light energy, such as leaf area density and distributions, tree density, local topography, and solar geometry (Walcroft et al., 2005). Light acclimation has been shown to affect respiration rates in white spruce dramatically, with higher rates of respiration in higher light environments (Awada & Redmann, 2000). These indirect environmental effects on leaf respiration may be particularly important at the extremes of white spruce distribution, given that there are intra-canopy gradients in light absorption in the dense canopy at the southern limit of the range that do not exist at the more open northern limit (Schmiege et al., *pers com*). Considering the vast potential physical, environmental and morphological differences across the white spruce range, it is important to consider the possible effects of canopy position on foliar respiration and its response to temperature.

The goal of this research was to gain a better understanding of how respiration varies between the two extremes of the white spruce species range and with canopy position at each of those extremes. Using our recent, detailed assessment of white spruce respiratory physiology at the Alaskan FTE (Griffin et al., 2021), we implement identical techniques to compare this to the physiological function of spruce growing more than 5000 km away at the opposite end of the species distribution. We test four research hypotheses. First, that leaf respiration measured at 25 °C (*R*25) will increase with latitude (Atkin et al., 2015). We extend this to examine the overall temperature response of respiration at the two range limits. Second, while our previous study did not find canopy position differences in respiration at the northern FTE site (Griffin et al., 2021), we hypothesize that at our southern site, upper canopy leaves will have higher respiration rates and be less responsive to temperature than lower canopy leaves. Third, we hypothesize that southern but not northern white spruce respiration will be similar to the average for the Needle-leaved Evergreen (NLEv) plant functional type (PFT) to which they belong. To quantify white spruce respiratory temperature response and make these comparisons, we use two models, the global polynomial model of Heskel et al. (2016) and the thermodynamic model of Liang et al. (2017). Fourth, we hypothesize that trees from the southern edge of the species range will have a higher temperature tolerance than trees from the northern edge of the distribution. To test this, we quantify *T*max, the leaf temperature at which the maximum rate of respiration is reached, from the respiratory temperature response curves. Finally, we examine the general relationships between respiratory traits and leaf traits, and test the prediction that leaf nitrogen will be a strong predictor of respiration. The two specific locations of this study are Black Rock Forest, located in the Hudson highlands of New York, and the FTE in north central Alaska, USA (referred to as the “southern location” and “northern location” respectively, Figure 1). Our southern site is at the same location as the species range study described above (Patterson et al., 2018) and represents an extreme southerly location for white spruce. By quantifying these variables at the extremes of the species distribution, we characterize the mechanistic contribution of respiration to the current and potential future distribution of white spruce.

## Materials and Methods

### Site Descriptions and Leaf Material

This research was conducted at two sites representing the opposite ends of the species range, Black Rock Forest (BRF), New York, USA (41°24’ 03.91” N latitude, 74°01’28.49” W longitude – southern location) and the Forest Tundra Ecotone (FTE), AK USA (67°59’ 40.92” N latitude, 149°45’15.84” W longitude – northern location). The sites are more fully described elsewhere (Eitel et al., 2019, Patterson et al., 2018). The average annual precipitation and temperature of these two sites are 1285 mm and 10.9 °C at the southern location (Arguez et al., 2010) and 485.4 mm and -8.12 °C at the northern location (Eitel et al., 2019). The northern location is spruce dominated evergreen forest (Eitel et al., 2019), while the southern location is a northern temperate deciduous forest that is oak dominated (Patterson et al., 2018; Schuster et al., 2008). The southern location is to the south of the natural species range distribution presented by Little (2004). White spruce was probably introduced into BRF as nursery stock for forestry trials in the early 1930’s but has since expanded naturally. The trees used for this study were naturally seeded on the forest edge along Continental Road, a dirt trail established during the Revolutionary War to facilitate troop travel between West Point and the encampment of the Continental army in New Windsor. While the trail represents a break in the forest canopy it gets limited use and has only a minor impact on the canopy-dominant study trees located at least 15 m from the road. Photographs of trees from the two sampling locations are shown in Figure S1.

Leaves used for the respiration measurements from the southern site were collected in late June and early July of 2018. Samples were taken from south facing branches of six study trees, and were selected from both upper (1 m below the apical meristem) and lower (1.37 m from the ground) canopy positions. As in our FTE study (Griffin et al., 2021), the terminal portions of several branches were cut with sharp pruners and the removed portion of the stem was immediately wrapped with wet paper towels, sealed in a plastic bag with ample air and placed in cooler where they could be kept dark and transported to the lab. The top of the canopy was accessed with an articulating boom lift. Once returned to the lab, the stem pieces were recut underwater and then placed in a beaker containing enough water to submerge the cut end until analyzed, typically within 8 (but no more than 24) hours. Leaves from the northern site were a subset of data used by Griffin et al., (2021) and thus are described fully in that study. The motivation of Griffin et al., (2021) was to assess the effect of tree size (saplings vs. trees) and test current hypotheses about the location of treeline. For the current study, we focus on fully developed trees growing at the northern and southern range extremes. Thus, the Griffin et al. (2021) dataset was trimmed to exclude all saplings (stems < 10cm DBH), leaving 18 individual trees (≥ 10 cm DBH) and re-analyzed for the present study to complement new data presented from the southern site. No part of the analysis presented here was included in the earlier publication.

### Respiration Temperature Response Curves

The techniques employed here were identical to those used in our treeline study in order to facilitate direct comparison and methods are described more fully in Griffin et al. (2021). Briefly, CO2 exchange rates were measured on needles that were carefully removed from the stems, weighed to determine the initial fresh mass (g), photographed to determine projected surface area and placed in a fine nylon mesh bag. The mesh bag containing the leaves was placed inside a custom-made cuvette (Patterson et al. 2018, Li et al. 2019, Schmiege et al. 2021) with computer controlled thermoelectrical cooling (CP-121 Thermoelectric Peltier Cooling Unit, TE Technology, Traverse City, MI USA). The custom cuvette was interfaced with a portable photosynthesis system (Li-6400XT, LiCor Lincoln, Nebraska USA) which recorded all gas exchange and environmental parameters every 20 seconds.

After equilibrating the system to 5°C, the mesh bag holding the leaves was sealed inside. Once stability was reached the instrument was zeroed and the response curve was measured as described in Heskel et al. (2016), O’Sullivan et al. (2013), and Schmiege et al. (2021). During measurements the flow rate through the cuvette was set to 500 ml min^-1^ and the CO2 concentration to 400 ppm. The air was dried using a Li-6400XT desiccant column and then transpiration was allowed to humidify the cuvette. During the measurement the cuvette temperature was ramped up from 5 to 65 °C at a constant rate of 1 °C min^-1^.

### Leaf Traits

Leaves were photographed with a known scale and *ImageJ* software was used to determine their projected area (Schneider et al., 2012). The leaves were then dried at 65°C for a minimum of 48 hours and again weighed to determine leaf dry mass (g). Specific leaf area (SLA cm^2^ g^-1^) was calculated and used to determine mass-based respiratory fluxes from the area-based fluxes. Leaf water content (%) and leaf dry matter content (LDMC -mg dry mass g^-1^ fresh mass) were calculated from the fresh and dry masses (Vaieretti et al., 2007). Leaf carbon, nitrogen and the stable isotope ratios of these elements were determined on Thermo Scientific Delta V+ IsoLink IRMS System (Thermo Fisher Scientific Inc., Waltham, MA, USA) at the Stable Isotope Lab of Lamont-Doherty Earth Observatory (Palisades, NY).

### Data Analyses

The respiration temperature response curves were analyzed as in Heskel et al. (2016) by fitting a second-order polynomial model to the log transformed respiration rates between 10 and 45 °C.

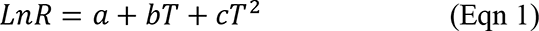

Where *a* represents the basal respiration rate (y-intercept), while *b* and *c* describe the slope and curvature of the response (Heskel et al. 2016). The respiration rate at a common temperature of 25 °C (*R*25) was also calculated and the temperature where the maximum respiration rate was reached (*T*max) was recorded. The modeled temperature response was also used to quantify the possible effect of warming on respiration rates (in the absence of thermal acclimation – *see justification below*), based on the current growing season average temperatures for the two sites and projections for end of the century warming (US Federal Government, 2021).

In addition, the thermodynamic underpinnings of the respiratory temperature response were explored using the model developed by Liang et al., (2017) based on macromolecular rate theory (MMRT). This model, which itself is based on transition state theory (TST) for enzyme-catalyzed kinetics, not only provides a thermodynamic explanation for the observed temperature response of *R*, but has also been extensively cross checked with the global database used by Heskel et al., (2016) to develop the simple log polynomial model above (Eqn 1). The model provides the following equivlencies to the coefficients in Eqn 1:

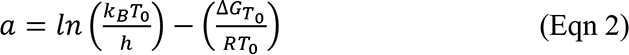

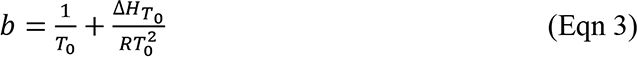

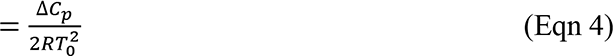

Where *k*B = the Boltzman constant, *h* = Plank’s constant, *R* = ideal gas constant, *T*0 = reference temperature. Δ*G*_*To*_ = Gibbs free energy between the ground state and transition state at the reference temperature and thus accounts for changes in both the change in bond energies (Δ*H*_*To*_) and the temperature dependent change in entropy (*TΔS*). Δ*C*_*p*_ representes the change in heat capacity during the reaction.

Respiration vs. temperature curves were fit to the polynomial model of Heskel et al., (2016) using R v. 3.6.3 (R Core Team, 2020). Due to the unequal sample size (n BRF = 6; n FTE = 18), a one-way ANOVA was used to test our first hypothesis regarding respiration across the species range. To test our second hypothesis regarding the main effects of canopy position on respiration *within* each of the two sites, paired t-tests were used to compare each of the model parameters *a*, *b* and *c* (see Eqn 1), and model predictions of *R25*. All traits were transformed as necessary to fulfil assumptions of normality. To test our third hypothesis, we compared our model estimates with those of Heskel et al. (2016) by means of an independent sample t-test based on the mean, confidence intervals and sample size for the NLEv PFT (as reported in their Table 1). Our fourth hypothesis regarding the *<u>T</u>*max of respiration was tested similarly to hypotheses 1 and 2 described above. Finally, regression equations were used to assess the general relationships between leaf traits and leaf respiration and, in particular, the ability of leaf N (per unit leaf area) to predict the rate of respiration (Atkin et al., 2015). All data analysis other than the initial R/T curve fitting was done in Excel (version 16.51 for Mac, Microsoft, Redmond, Washington, U.S.A.) with both the Solver and *RealStatistics* (Release 7.6, Zaiontz 2021) add-ins.

**Table 1.**
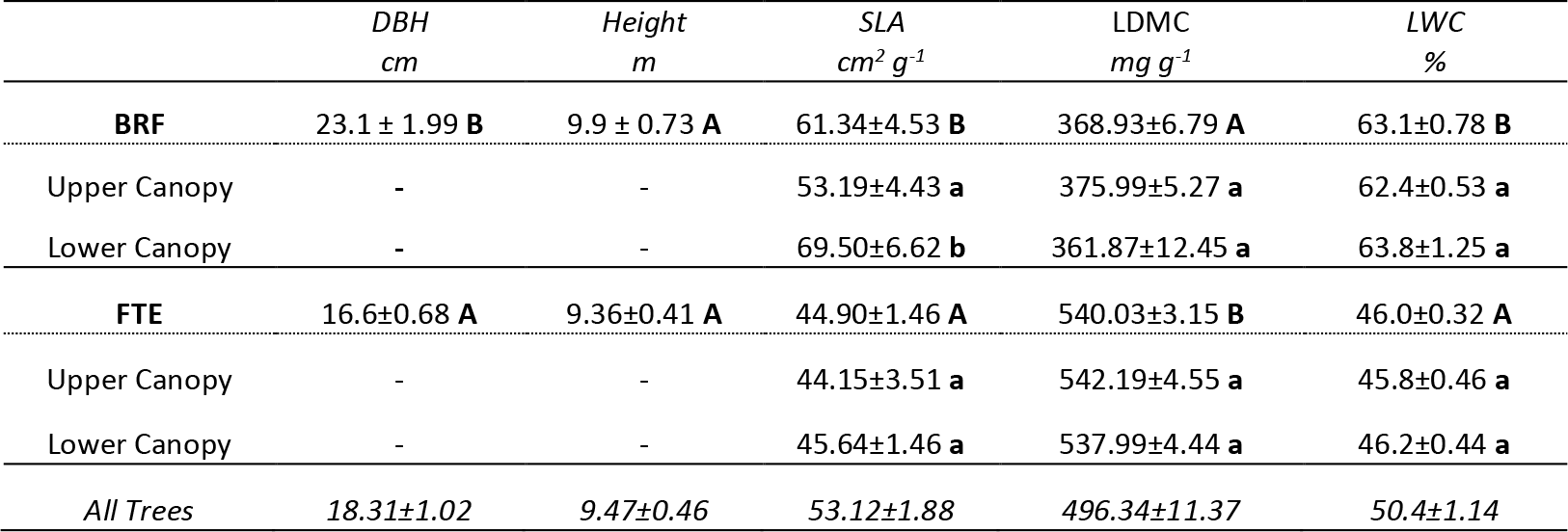
Top of canopy leaf and bottom of canopy tree and leaf characteristics of Picea glauca growing at either the southern edge of the species range limit in Black Rock Forest (BRF), New York, USA, or at the northern edge at the Forest Tundra Ecotone (FTE) in northcentral Alaska, USA. DBH = diameter at breast height (1.37m), SLA = specific leaf area, LDMC = leaf dry matter content (g dry mass g^-1^ fresh mass), LWC = leaf water content and N = leaf nitrogen %. Lowercase letters following the canopy position means (within a site) and uppercase letters following the location means (across the canopy positions) denote statistically significant differences (p<0.05). n = 6-47, all values mean±1 standard error.

|  | DBH<br>cm | Height<br>m | SLA<br>cm<sup>2</sup> g<sup>-1</sup> | LDMC<br>mg g<sup>-1</sup> | LWC<br>% |
| --- | --- | --- | --- | --- | --- |
| **BRF** | 23.1 ± 1.99 **B** | 9.9 ± 0.73 **A** | 61.34±4.53 **B** | 368.93±6.79 **A** | 63.1±0.78 **B** |
| Upper Canopy | - | - | 53.19±4.43 **a** | 375.99±5.27 **a** | 62.4±0.53 **a** |
| Lower Canopy | - | - | 69.50±6.62 **b** | 361.87±12.45 **a** | 63.8±1.25 **a** |
| **FTE** | 16.6±0.68 **A** | 9.36±0.41 **A** | 44.90±1.46 **A** | 540.03±3.15 **B** | 46.0±0.32 **A** |
| Upper Canopy | - | - | 44.15±3.51 **a** | 542.19±4.55 **a** | 45.8±0.46 **a** |
| Lower Canopy | - | - | 45.64±1.46 **a** | 537.99±4.44 **a** | 46.2±0.44 **a** |
| *All Trees* | 18.31±1.02 | 9.47±0.46 | 53.12±1.88 | 496.34±11.37 | 50.4±1.14 |

## Results

### Tree and leaf traits

The average tree in this study was 18.3±1.0 cm in DBH (1.37 m from the ground) and 9.5 ±0.46 m tall (Table 1). Trees from the southern location tended to be larger in diameter (23.1±2.0 vs. 16.6±0.7 cm), but similar in height (9.9 ±0.73 vs. 9.36±0.41 m) compared to the northern location trees. The leaf dry matter content (LDMC) and leaf water content (LWC) did not differ by canopy position at either location, but LDMC was 32% lower and LWC was 37% higher at the southern location compared to the northern location (Table 1). The specific leaf area (SLA) of the upper canopy leaves was 23% lower than that of the lower canopy leaves at the southern location (Table 1). Leaf C did not vary between the two sites, but leaf N concentration ([N]) was 36% lower at the northern compared to the southern location and thus the C/N ratio was 56% higher at the FTE (Table 2). Canopy position did not affect either the leaf C or C/N ratio at either site (Table 2). The lower canopy leaves of the southern location had the most negative δ^13^C and were significantly different than the upper canopy leaves at this same site. As a result, leaves from the southern location had significantly more negative δ^13^C values than leaves from the northern site (Table 2). Leaf δ^15^N also differed significantly between the two locations with the northern location being the more negative of the two (Table 2). On a leaf area basis, leaf nitrogen was 70% higher at the northern location than the southern location (Table 2). Canopy position did not affect leaf nitrogen per unit area at either location.

**Table 2.** Top of canopy leaf and bottom of canopy leaf chemistry of Picea glauca growing at either the south-eastern edge of the species range limit in Black Rock Forest (BRF), New York, USA, or at the northern edge at the Forest Tundra Ecotone (FTE) in northcentral Alaska, USA. Lowercase letters following the canopy position means (within a site) and uppercase letters following the location means (across the canopy positions) denote statistically significant differences (p<0.05). n = 6-47, all values mean±1 standard error.

| | C<br>% | N<br>% | C/N | $\delta^{13}\text{C}$<br>‰ | $\delta^{15}\text{N}$<br>‰ | N<br>g m<sup>-2</sup> |
| --- | --- | --- | --- | --- | --- | --- |
| **BRF** | 45.49 ± 0.36 **A** | 1.29 ± 0.07 **B** | 36.41 ± 1.84 **B** | -29.11 ± 0.29 **B** | -0.10 ± 0.21 **B** | 2.20 ± 0.17 **A** |
| Upper Canopy | 45.63 ± 0.42 **a** | 1.28 ± 0.08 **a** | 36.26 ± 2.12 **a** | -28.34 ± 0.38 **a** | 0.13 ± 0.25 **a** | 2.47 ± 0.21 **a** |
| Lower Canopy | 45.34 ± 0.63 **a** | 1.29 ± 0.11 **a** | 36.57 ± 3.23 **a** | -29.76 ± 0.24 **b** | -0.34 ± 0.33 **a** | 1.93 ± 0.25 **a** |
| **FTE** | 46.09 ± 0.22 **A** | 0.82 ± 0.02 **A** | 56.96 ± 1.24 **A** | -28.51 ± 0.14 **A** | -7.08 ± 0.28 **A** | 3.78 ± 0.10 **B** |
| Upper Canopy | 46.07 ± 0.32 **a** | 0.82 ± 0.02 **a** | 57.07 ± 1.79 **a** | -28.43 ± 0.20 **a** | -7.12 ± 0.43 **a** | 3.81 ± 0.13 **a** |
| Lower Canopy | 46.11 ± 0.31 **a** | 0.82 ± 0.03 **a** | 56.85 ± 1.76 **a** | -28.59 ± 0.21 **a** | -7.03 ± 0.38 **a** | 3.76 ± 0.16 **a** |
| *All Trees* | 45.94 ± 0.19 | 0.94 ± 0.04 | 51.82 ± 1.65 | -28.66 ± 0.13 | -5.33 ± 0.49 | 3.38 ± 0.13 |

### Respiration temperature response curves

In all trees, respiration increased exponentially between 5 and 45 °C, then slowed briefly before increasing rapidly to a maximum rate (*R*max) defining the *T*max (Figure 2a). Measured respiration rates between 10 and 45 °C were used to model the ecologically relevant response (Figure 2b). The global polynomial model of Heskel et al. (2016) fit all log normal respiration temperature curves with an r^2^ ≥ 0.99 (Figure 3). Overall, the three model coefficients averaged -1.82±0.12, 0.090±0.003 and -0.00030±0.0001 (*a*, *b* and *c* respectively, mean ± SEM, Table 2). At the southern location, leaves from the top of the canopy had a 24% lower intercept (coefficient *a*) than leaves from the bottom of the canopy (p=0.04, Table 3). Canopy position did not affect the model coefficients at the northern location. The temperature response of white spruce growing at the southern location was quite similar to the average model coefficients for the plant functional type to which this species belongs, needle-leaved evergreen (NLEv, Heskel et al. 2016), and only the curvature (*c*) of the southern location upper canopy leaves was statistically different from the average NLEv response (Figure 3).

**Figure 2.**
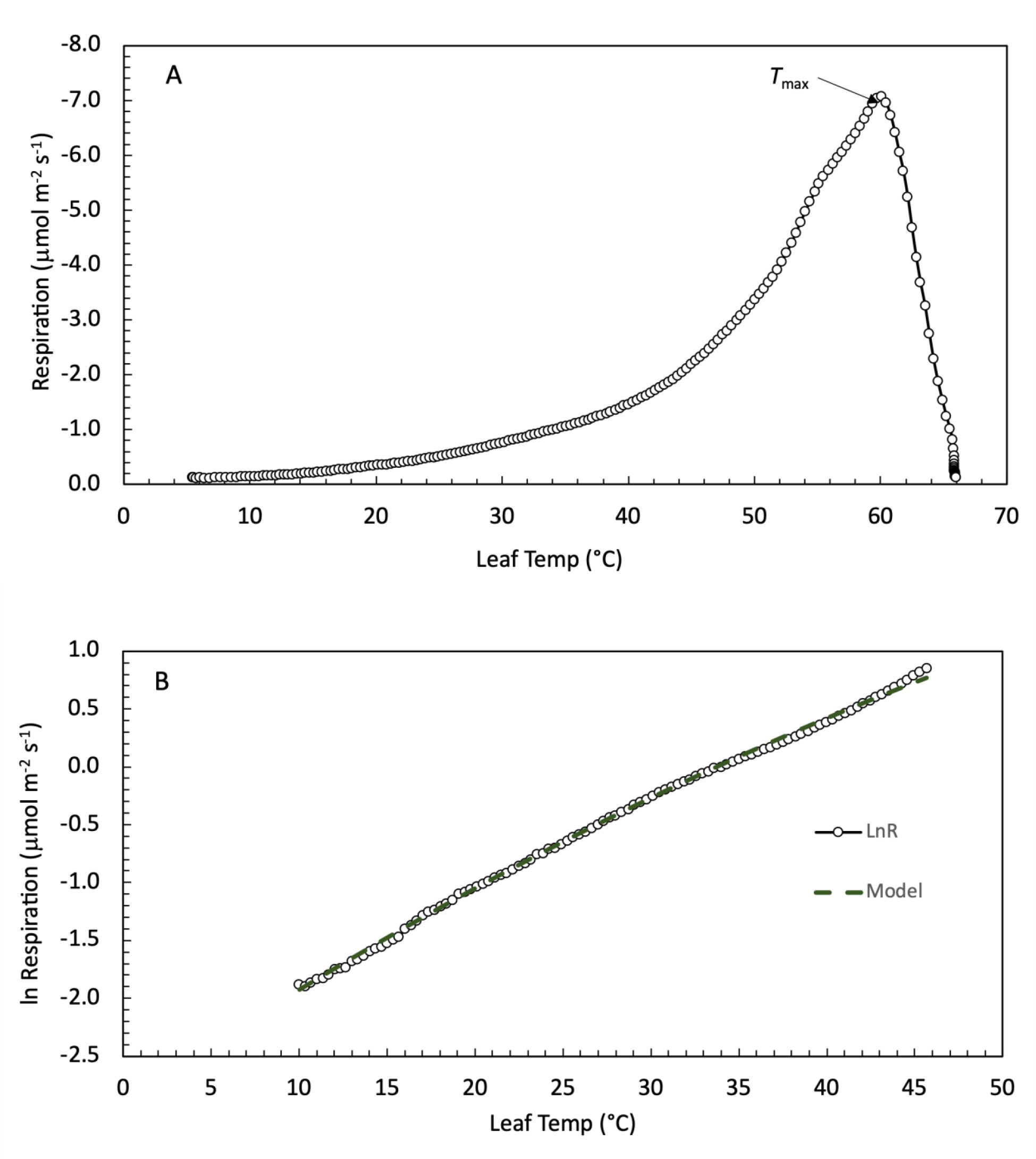
Leaf respiration as a function of temperature in *Picea glauca* measured at the southern end of the species distribution in Black Rock Forest, New York, USA. Panel A (top) is an example high-resolution temperature response curve measured from 5 to 65 °C. Air temperature was heated at a rate of 1 °C min^-1^ while the rate of CO2 release and other gas-exchange parameters were recorded every 20 seconds. Panel B (bottom) is the log of measured respiration rate and model fit (*lnR = a + bT + cT^2^* – see text for full description) between 10 and 45 °C for the same sample.

**Figure 3.**
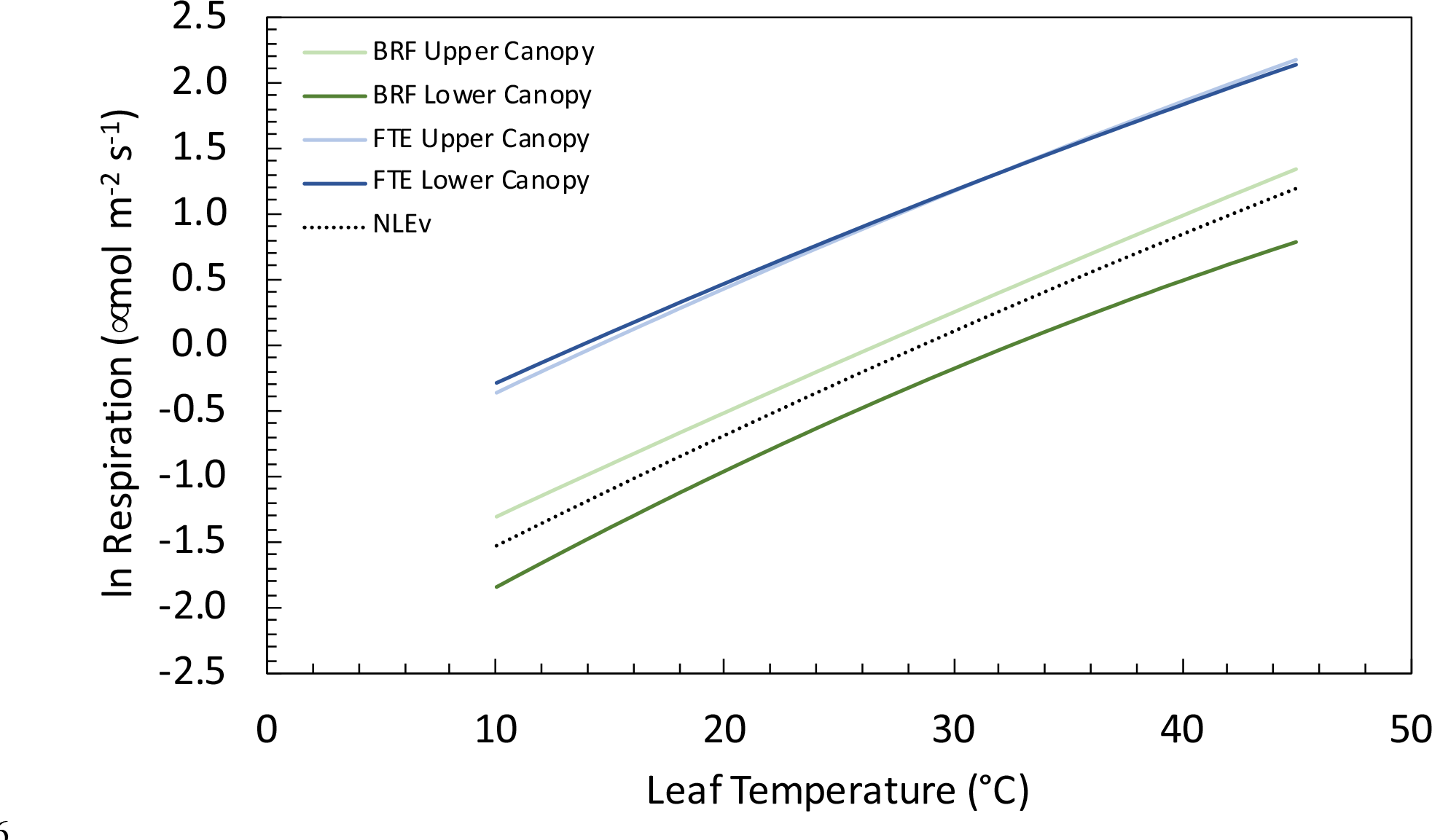
Average model results (log of leaf respiration) for the bottom of canopy (dark green and blue lines) and top of the canopy (light green and blue lines), of *Picea glauca* trees growing at either the southern edge of the species range (green lines) in Black Rock Forest (BRF), New York, USA, or the northern edge of the species range limit (blue lines) in the Forest Tundra Ecotone (FTE) in northcentral Alaska, USA. Line presents the mean response (n=6 (BRF) or 18 (FTE). Also shown is the average response for Needle-Leaved Evergreen species (NLEv) from the global survey of Heskel et al (2016) (black dotted line). Model parameters are presented in Table 2.

**Table 3.** Polynomial model parameters (Heskel et al., 2016) fit to the measured high-resolution leaf respiration temperature response curves collected from either the top of canopy leaves or bottom of canopy leaves of *Picea glauca* trees growing at either the southern edge of the species range limit in Black Rock Forest (BRF), New York, USA, or the northern edge of the species range limit in the Forest Tundra Ecotone (FTE) in northcentral Alaska, USA. Lowercase letters following the canopy position means and uppercase letters following the location means denote statistically significant differences (p<0.05). n = 6-47, all values mean±1 standard error.

|  | *a* | *b* | *c* |
| --- | --- | --- | --- |
| **BRF** | -2.48±0.56 **B** | 0.094±0.009 **A** | -0.00034±0.0001 **A** |
| Upper Canopy | -2.166±0.105 **a** | 0.084±0.006 **a** | -0.00014±0.0001 **a** |
| Lower Canopy | -2.829±0.371 **b** | 0.104±0.017 **a** | -0.00053±0.0003 **a** |
| **FTE** | -1.15±0.09 **A** | 0.086±0.003 **A** | -0.00027±0.0001 **A** |
| Upper Canopy | -1.20±0.29 **a** | 0.087±0.004 **a** | -0.00028±0.0001 **a** |
| Lower Canopy | -1.10±0.27 **a** | 0.084±0.005 **a** | -0.00026±0.0003 **a** |
| *All Trees* | 1.82±0.12 | 0.090±0.003 | -0.00030±0.0001 |

However, the southern location measurements from both canopy positions were significantly different when compared to their counterparts from the northern location (Figure 3). These differences include lower intercepts for both the BRF canopy positions, as well slight differences in both the slope and curvature of the upper canopy leaves. The response of respiration to temperature at the northern site was significantly different from the NLEv PFT response (higher *a*). The Gibbs free energy released during the transition from the ground state to the transition state of the respiratory process was 4% higher from leaves at the southern location compared to the northern location (Table 4).

**Table 4.** Macromolecular rate theory (MMRT) model parameters (Liang et al 2017) calculated from the measured high-resolution leaf respiration temperature response curves collected from either the top of canopy leaves or bottom of canopy leaves of Picea glauca trees growing at either the southern edge of the species range limit in Black Rock Forest (BRF), New York, USA, or the northern edge of the species range limit in the Forest Tundra Ecotone (FTE) in northcentral Alaska, USA. **ΔGT** = the difference in Gibbs free energy between the ground state and transition state, **ΔHT** = the change in enthalpy and **ΔCP** = change in heat capacity. Lowercase letters following the canopy position means and uppercase letters following the location means denote statistically significant differences (p<0.05). n = 6 - 47, all values mean±1 standard error.

| | $\Delta G_T$<br>kJ mol<sup>-1</sup> | $\Delta H_T$<br>kJ mol<sup>-1</sup> | $\Delta C_p$<br>kJ mol<sup>-1</sup> K<sup>-1</sup> |
| --- | --- | --- | --- |
| **BRF** | $0.79 \pm 0.005$ B | $67.04 \pm 6.824$ A | $-0.50 \pm 0.228$ A |
| Upper Canopy | $0.78 \pm 0.003$ a | $59.39 \pm 4.163$ a | $-0.21 \pm 0.141$ a |
| Lower Canopy | $0.80 \pm 0.009$ a | $74.69 \pm 12.812$ a | $-0.79 \pm 0.419$ a |
| **FTE** | $0.76 \pm 0.002$ A | $60.84 \pm 2.11$ A | $-0.40 \pm 0.066$ A |
| Upper Canopy | $0.76 \pm 0.003$ a | $62.12 \pm 2.611$ a | $-0.44 \pm 0.080$ a |
| Lower Canopy | $0.76 \pm 0.003$ a | $59.62 \pm 3.330$ a | $-0.37 \pm 0.106$ a |
| *All Trees* | $0.77 \pm 0.003$ | $62.42 \pm 2.336$ | $-0.43 \pm 0.075$ |

### Respiration at a common temperature & at Rmax

Across all samples at the southern location, the average rate of respiration at a common temperature of 25 °C was 0.75 ± 0.08 μmol CO2 m^-2^ leaf area s^-1^, which is 68% lower than the average rate at the northern location (2.35±0.88 μmol CO2 m^-2^ leaf area s^-1^). The rate differed significantly by canopy position at the southern, but not the northern location. *R*25 was 48% higher in upper canopy leaves than in lower canopy leaves (0.90 ± 0.09 μmol m^-2^ s^-1^ vs. 0.61 ± 0.10 μmol m^-2^ s^-1^) from the southern location (Figure 4a). Due to significantly lower SLA of the southern location upper canopy leaves compared to the lower canopy leaves, there were no significant differences in the mass based *R*25 (Figure 4b). Per gram of leaf N, the respiration rates at 25 °C did not differ by canopy position at either site, but overall the northern location had rates that were 87% higher than those of the southern location (0.34 ± 0.019 vs. 0.64 ± 0.034 μmol g N^-1^ s^-1^, Figure 4c). Leaves from the southern location, at both canopy positions, continued to respire until leaf temperature reached 58.5 ± 0.5 °C (*T*max) before quickly decreasing as the leaves died. This was a slightly higher, but statistically similar temperature to the *T*max of northern location trees (57.6±0.36 °C). However, at the southern location the maximum rate of respiration (*R*max) of the upper canopy leaves at *T*max (7.43 ± 0.78 μmol m^-2^ s^-1^) was 40% higher than the *R*max of the lower canopy leaves (5.29 ± 0.66 μmol m^-2^ s^- 1^), and the southern location average *R*max was 59% lower than the northern location average, a difference of more than 9.3 μmol CO2 m^-2^ leaf area s^-1^ (Figure 4d). All of the major respiratory parameters (*R*25, *a* and *R*max) were significantly related to leaf N (mg N m^-2^). *R*25 at the northern location was nearly twice the *GlobResp* average and more than double the *GlobResp* gymnosperm average (Figure 5). There was a significant negative relationship between *R*25 per unit leaf area and the average growing season temperature across published respiration rates for this species (Figure 6, Table S1). Estimated *in situ* rates of respiration at the average growing season (i.e. June, July, August) temperature were significantly higher at the northern location than at the southern location (indicated as solid vertical arrows on Figure 7), as are the estimated rates for the projected end of century temperatures (indicated by dotted vertical arrows on Figure 7). These projections do not account for thermal acclimation. Benomar et al. (2018) report that while basal respiration is linearly related to growing season temperature, the shape of the temperature response (Q10) does not change across an experimental gradient varying in temperature.

**Figure 4.**
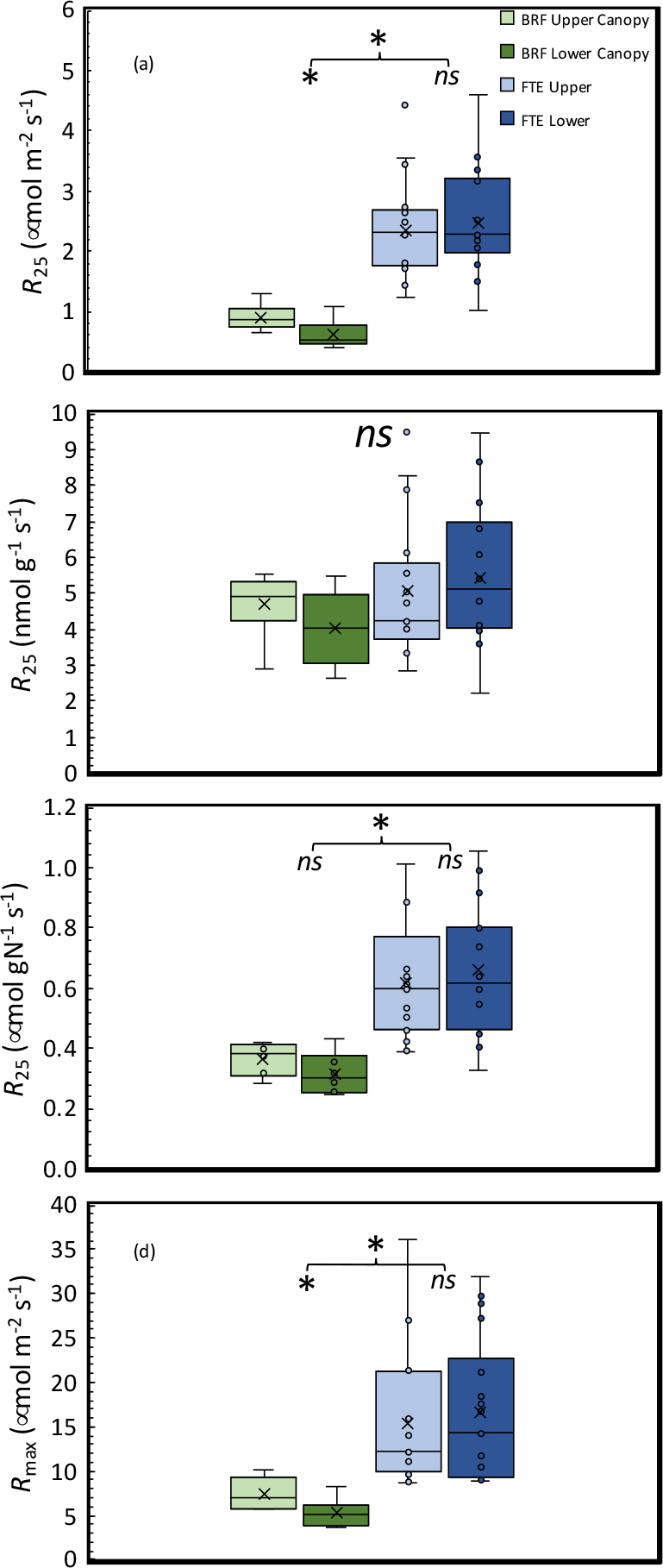
Respiration of white spruce (*<u>Picea glauca</u>*) foliage measured from the upper (light green) or lower (dark green) canopy in the southern location (Black Rock Forest (BRF), New York, USA), and the upper (light blue) or lower (dark blue) canopy from the northern location (Forest Tundra Ecotone (FTE), Alaska, USA). Respiration was measured at a leaf temperature of 25 °C on a projected leaf area basis (panel a), leaf mass basis (panel b), leaf nitrogen basis (panel c), as well as the maximum rate of respiration from a temperature response curve, expressed on a leaf area basis (panel d). The middle line of the box represents the median, the x represents the mean. The box is drawn between the second and third quartiles and the whiskers extend to the first and fourth quartiles. * = statistical significance (p<0.05). *ns* = not significant. n = 6 (BRF), 18 (FTE)

**Figure 5.**
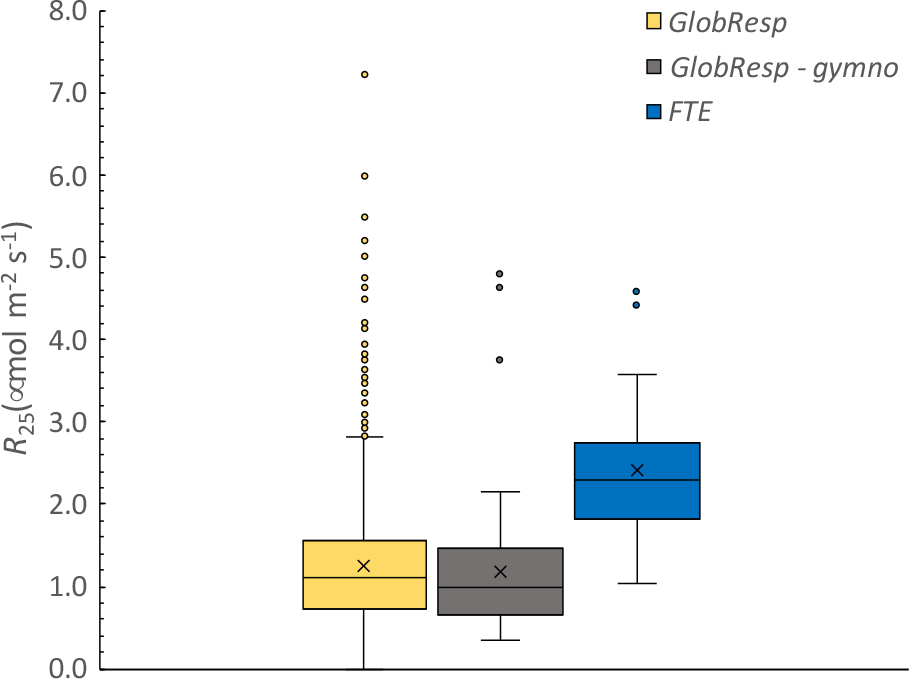
Comparison of leaf respiration of white spruce (*<u>Picea glauca</u>*) foliage measured at the Forest Tundra Ecotone (FTE, Alaska, USA – blue) with all measurements from the *GlobResp* database (Atkin et al., 2015 - yellow) and the gymnosperm measurements from the same database (brown). All respiration rates were measured at a leaf temperature of 25 °C and are expressed on a projected leaf area basis. The middle line of each box represents the median; the x represents the mean. The box is drawn between the second and third quartiles and the whiskers extend to first and fourth quartiles. N = 1114 (*GlobResp*), 57 (*GlobResp* Gymno), 18 (FTE).

**Figure 6.**
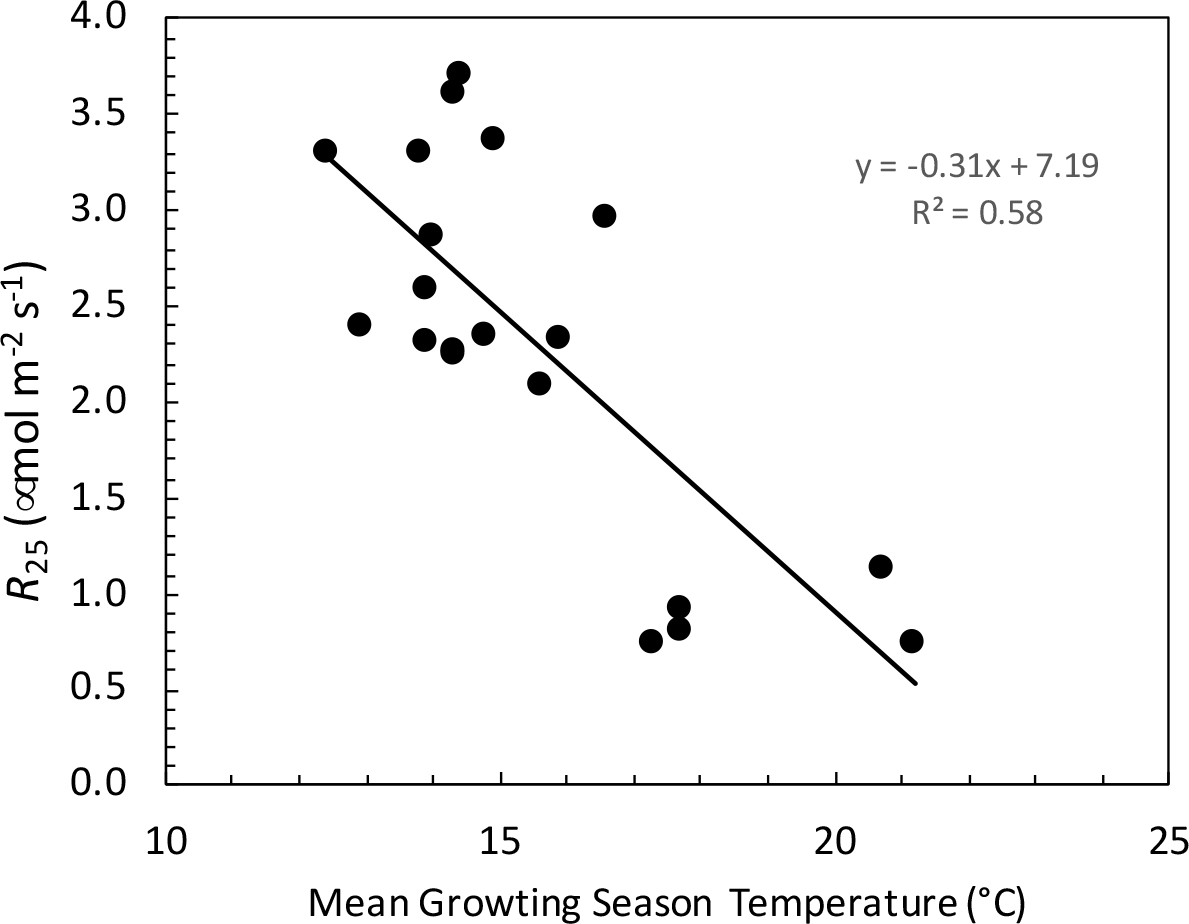
Summary of the relationship between published White Spruce (*<u>Picea glauca</u>*) *R*25 (leaf respiration measured at 25 °C), and the average growing season temperature of the study locations. *p* = 0.00001. Data sources can be found in the supplemental information.

**Figure 7.**
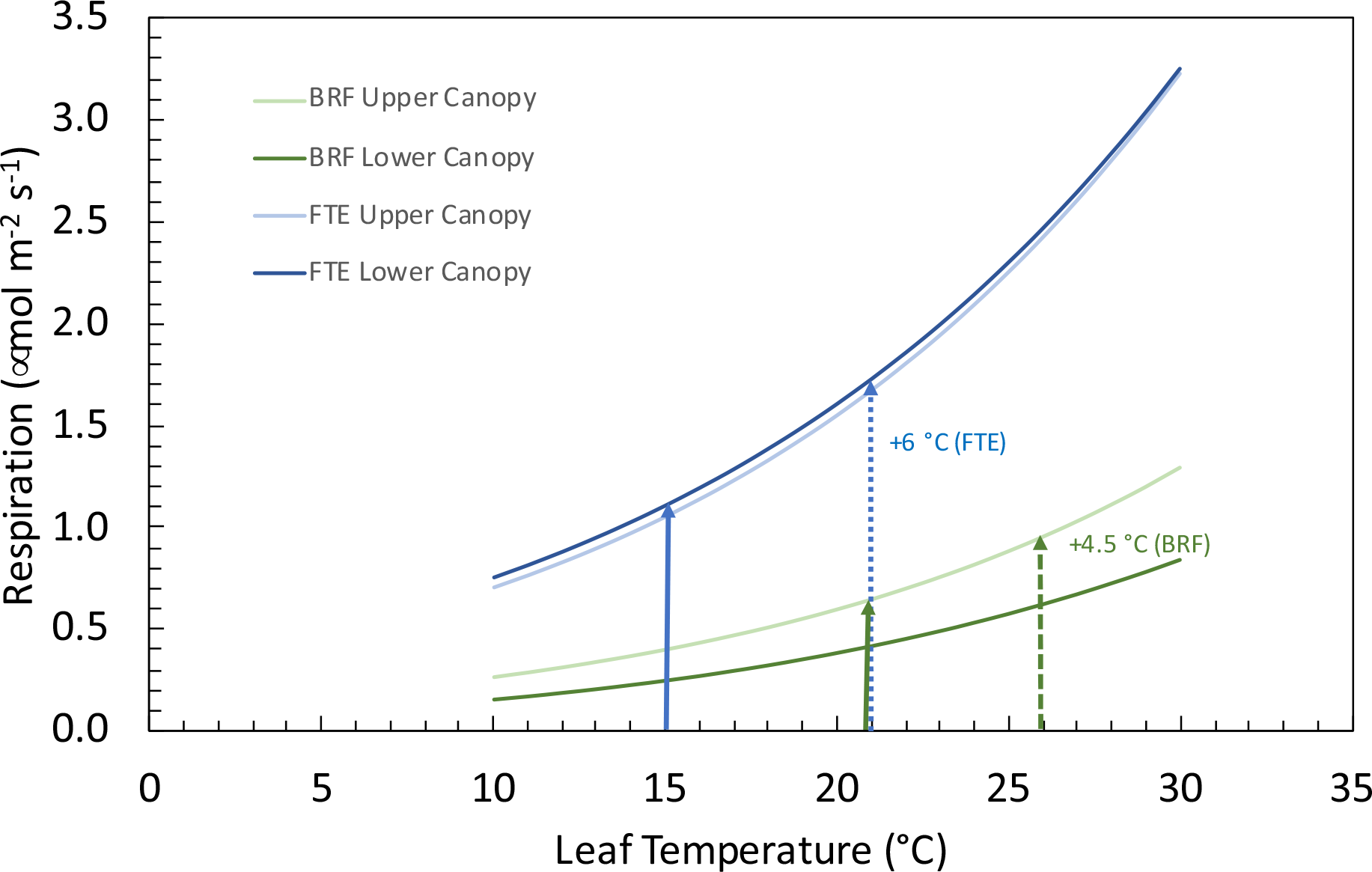
Modeled rates of white spruce (*<u>Picea glauca</u>*) foliage respiration vs. leaf temperature from two canopy positions at either the northern or southern range limits of the species. Lines display the average modeled response from the upper (light green) or lower (dark green) canopy from the southern location (Black Rock Forest (BRF), New York, USA, n= 6), and the upper (light blue) or lower (dark blue) canopy from the northern location (Forest Tundra Ecotone (FTE), Alaska, USA, n = 18). Vertical arrows represent the average June, July, August temperatures from each of the field sites (solid blue = FTE, solid green = BRF) and the projected end of century increases (dashed blue = FTE projection, dashed green = BRF projections, see text for more information).

## Discussion

We found dramatic differences in white spruce leaf respiration between trees at the limits of the species range. Our two locations are separated by more than 5000 km, from New York State at the south-eastern edge of the distribution, to the Forest Tundra Ecotone in arctic, north-central Alaska. The differences are large, and likely represent a significant physiological response by this species to differences in ambient environmental conditions. The rates of respiration at the northern range limit of the species distribution are nearly twice as high (when measured at a common reference temperature) as the global average and more than twice as high as the average gymnosperm according to a global database of leaf respiration rates (*GlobResp*, Atkin et al., 2015). At the limits of the vast range of this species, many environmental differences exist, including in day length, air temperature, soil temperature, soil moisture, vapor pressure deficit (VPD), pollutant exposure and edaphic factors, all of which are moderated differently by the distinctive forest canopy structures of the sites. Of the above environmental factors, the response of white spruce foliage to air temperature is of particular interest for three reasons: 1) temperature is known to have strong effects on the rate of respiration (Heskel et al., 2016); 2) temperature is increasing rapidly and unevenly across the species range (warming is more rapid at high latitudes (Cohen et al., 2014; Huang et al., 2017) and; 3) ecosystem models calculate the temperature response of respiration to scale over both time and space (Atkin et al., 2014; Heskel et al., 2016; Huntingford et al., 2017a). At the northern and southern range limits of white spruce, average growing season (June, July, and August) temperatures differ by nearly 6 °C, from approximately 21 °C at the southern location (US Federal Government, 2021) to slightly less than 15 °C at the northern location (Harris et al., 2020; Zepner et al., 2021). This 29% difference in average temperature is associated with a tripling of the respiration rate in white spruce growing at the FTE relative to the southern site when measured at a common temperature of 25 °C, and a doubling of the rate at the *in situ* average growing season temperature.

Combining our findings with published rates of leaf respiration in white spruce demonstrates a significant linear trend between decreasing growing season temperatures and increasing rates of respiration.

The much higher rates of respiration at the northern range limit of this species support our first research hypothesis and may help explain not only why white spruce is not found further to the north, but also why the boreal forest biome transitions to tundra at these high northern latitudes (Griffin et al., 2021). Plants acclimated to cold temperatures often have higher rates of respiration measured at a common temperature and a steeper respiratory temperature response (Atkin & Tjoelker, 2003; Körner, 1989; Reich & Oleksyn, 2004) as a means to maintain metabolic function in cold environments. A comparison of the mean growing season temperature and reported rates of leaf respiration across the white spruce range confirms this pattern in the species. The high respiratory rates, coupled with slow growth rates (Jensen et al., *pers com*) at the northern range limit suggest that northern trees experience large carbon losses that are likely related to high maintenance and other metabolic costs. These costs are not incurred at the same magnitude by white spruce growing at the southern edge of its distribution.

The response of respiration to temperature in trees at either end of the species range provides additional insights into the regulation of respiration and the root cause of the observed differences. First, we find differences between the two locations in the measured rates of respiration at both cold and warm temperatures. This finding is consistent with Type II acclimation (Atkin & Tjoelker, 2003) and suggests that there are differences in respiratory capacity, efficiency or perhaps mitochondrial numbers and structure (Atkin & Tjoelker, 2003; Klikoff, 1966; Kornfeld et al., 2013; Miroslavov & Kravkina, 1991; Patterson et al., 2018).

Second, the measured *Tmax* of white spruce needle respiration does not differ significantly between the two geographic extremes. This indicates that overall thermal tolerance is not plastic but a fixed species trait in white spruce (Heskel et al., 2014; O’Sullivan et al., 2013, 2017).

Third, we find differences in the temperature response of respiration across the species range, with northern white spruce also exhibiting a significantly different response to that of the needle leaved evergreen plant functional type. This suggests that the northern range limit is an extremely challenging environment for the species, and that the high respiration rates may be affecting leaf carbon balance and ultimately tree survival (Heskel et al., 2016; Huntingford et al., 2017b; Patterson et al., 2018). And fourth, despite the indication of a Type II acclimation response (Atkin & Tjoelker, 2003), the adjustments in respiratory physiology do not result in homeostasis with regards to maintaining observed rates of respiration at the local temperatures during the growing season. The lack of homeostasis may reflect the large difference in ambient temperatures that exist between the northern and southern range limits (Campbell et al., 2007). In summary, rates of respiration at the northern edge of the species range are high and respond strongly to temperature, most likely stemming from biochemical and physiological responses to the extreme environmental conditions of the Arctic.

The polynomial model we used to describe the temperature response of white spruce leaf respiration (Heskel et al., 2016) resulted in excellent predictions over the physiologically and ecologically relevant ranges of leaf temperatures and is useful for facilitating the biome and PFT comparisons above. The model of Liang et al. (2017), is derived from macromolecular rate theory, and provides a thermodynamic interpretation of these results. Term *a*, which varied by both location and canopy position, defines the rate of respiration as defined by variables such as substrate and moisture availability, and is thermodynamically related to the activation energy of the overall respiratory process (Liang et al., 2017). Our results are consistent with others who show that activation energy can be considered a temperature dependent feature of an ecosystem (Davidson & Janssens 2006; Lloyd & Taylor 1994; Liang et al., 2017). Furthermore, the *b* coefficient, which represents changing enthalpy (Δ*H*To) for the reaction at the reference temperature, is unaffected by either location or canopy position, and thus our examination of white spruce respiratory physiology is consistent with the broad patterns found when comparing biomes and PFT’s (Liang et al., 2017). Finally, we show that the heat capacity of the respiratory system (Δ*C*p) is similarly unaffected by either location or canopy position, and shows a convergent respiratory response to temperature. Once the differences in the basal rates are quantified, either model can accurately predict respiration of white spruce, despite the well-known temperature dependence of the activation energy of the overall reaction (Kruse & Adams 2008; Noguchi et al., 2015; O’Sullivan et al., 2013).

We gained further insights into the response of white spruce to local environmental conditions by examining leaf traits and the relationships between them and respiratory physiology. Needles from the northern edge of the species distribution tend to be thicker, or perhaps denser, than needles from the southern location, likely reflecting large differences in ambient temperatures (Atkin et al., 2006; Rosbakh et al., 2015). While leaf carbon concentrations were invariant at 45.94 ± 0.19 %, the higher carbon to nitrogen ratio of leaves at the northern range limit again suggests a relative increase in leaf structure at this site compared to the southern extreme. *R*25 per gram of leaf material is also statistically invariant at our two sites, further suggesting that the difference in respiration is not related to increased structure, but rather to the metabolic function of the leaves. Increasing leaf structure is potentially adaptive in the harsh environmental conditions at the arctic treeline where, in addition to cold temperatures, leaves must survive high winds and abrasion, prolonged winters and the possibility of large snow or ice loads while remaining metabolically active for many years. Overall, the variation in leaf structural characteristics were consistent with expected relationships with cold temperatures and canopy gradients (Niinemets et al., 2015).

Leaf nitrogen typically correlates negatively to temperature (Yin, 1993) and latitude (Körner 1989, Reich & Oleksyn 2004), and our findings show that these trends hold at the opposite ends of the white spruce species range. The low leaf nitrogen concentration at the northern compared to the southern range limit likely reflects the effects of environment on nitrogen biogeochemistry, which often results in nitrogen limitation at the northern location. For example, arctic temperatures, hydrology, permafrost dynamics and bedrock geology can all limit nitrogen availability (Nadelhoffer et al., 1991; Schimel et al., 2004). Furthermore, ecosystem properties such as limited nitrogen fixation, slow decomposition rates, rapid nitrogen uptake rates and microbial competition could further limit nitrogen availability at the northern range limit (Schimel & Chapin, 1996; Schimel & Bennett, 2004; Yano et al., 2010). While the precise mechanism or combination of mechanisms limiting nitrogen uptake at the northern location is unknown, it is interesting that the resulting low leaf nitrogen concentrations are offset by changes in SLA. The result is higher leaf nitrogen per unit leaf area at the northern range limit and positive correlations with foliar respiration rates. This relationship conforms to global observations (Reich et al., 2008) and facilitates the modeling and prediction of respiration (Atkin et al., 2015). Finally, per gram of nitrogen, leaf respiration is higher at the northern range limit, indicating higher maintenance costs at this site relative to the southern site.

The nitrogen isotope signature of the northern and southern sites differed significantly, which may reflect a change in nitrogen source, differences in within-plant fractionation, and/or mycorrhizae-to-plant fractionation (Pritchard & Guy 2005; Craine et al., 2009, 2015).

Experimentally, when spruce is grown with NO_3_ ^-^, the isotopic ratios become more negative than when grown with either NH4^+^ or NH4NO3 (Pritchard & Guy 2005). This, together with a positive correlation between the foliar nitrogen isotope ratio and tree growth in white spruce (Matsushima et al., 2012), may help explain the much lower (more negative) δ^15^N values at the northern range limit than at the southern range limit. However, spruce is ectomycorrhizal (Malloch & Malloch 1981), which tends to lower the isotope ratios relative to arbuscular and non-mycorrhizal species (Craine et al., 2009). Thus, a reduced rate of infection at the southern site could potentially contribute to the observed differences. Similarly, cold temperatures are associated with lower δ^15^N values (Craine et al., 2009). These factors are not mutually exclusive, nor strictly predictive, and thus attributing the large differences in observed δ^15^N to specific mechanisms is not possible without significantly more experimentation. Working in the same general location more than 25 years earlier, Schulze et al. (1994) reports a similar δ^15^N in this species (-7.7 ‰ vs. -7.1 ‰), suggesting the ecological differences in the treeline N cycle are long-lived. Here we connect these to respiratory activity and demonstrate that leaf nitrogen serves as a strong proxy for respiration (Atkin et al., 2015) as a result of morphological adjustments that maintain a high leaf nitrogen per unit leaf area despite limited nitrogen availability.

Finally, a small difference in carbon isotope fractionation was found between our two sites, suggesting that white spruce at the northern limit of the species range has slightly higher water use efficiency than trees at the southern limit. The lower canopy leaves at the southern range limit have the most negative carbon isotope ratios (-29.76 ± 0.24), perhaps reflecting canopy gradients in both light and VPD. Overall, these patterns in leaf traits outline the general adaptative strategy in white spruce necessary to survive harsh northern conditions, and implicate the local environmental conditions as directly and indirectly driving the respiratory physiology across the species range.

We found support for our second hypothesis, that needles from the top of the tree would have higher rates of respiration than needles from the bottom of the tree at the southern range limit where canopy complexity creates resource gradients. The nature of the tree crown and the forest canopy differs significantly at our two sites. The southern location trees are strongly conical and compete for above ground resources in a mostly closed canopy surrounded by deciduous hardwood species. The FTE trees, however, are more cylindrical and arranged in mostly open canopies composed of individual trees or small groups of trees with only limited competition for space and aboveground resources such as light. As a result, strong intra-canopy light gradients exist at the southern location but are much less pronounced at the northern location. Light, VPD, wind and air temperature microclimates could all be affected by these differences in canopy structure (Whitehead et al., 2001; Walcroft, et al., 2004), but light is the most plausible driver of the observed respiratory response (*similar to* Bond et al., 1999 or Niinemets et al., 2015 (photosynthesis)). The upper canopy leaves at the southern location receive full light and as a result have higher photosynthetic rates than the lower canopy leaves which experience self-shading and generally lower ambient light conditions (Schmiege et al., *pers com).* Higher photosynthetic rates and more rapid growth in the upper canopy require higher respiration to support the ensuing growth and maintenance costs (Bond et al., 1999; Griffin et al., 2001, 2002; Tissue et al., 2002; Whitehead et al., 2004, Xu and Griffin 2006; Weerasinghe et al., 2014; Araki et al., 2017). By contrast, the combination of the open canopy, low sun angles and narrow unbranching crown morphology of white spruce at the northern location has the effect of homogenizing the light environment, and thus equalizing both photosynthetic capacities (see Niinemets et al., 2015) and the rates of respiration throughout the canopy. Clearly spruce trees at either end of the species distribution must dynamically respond and adapt to different environmental conditions over multiple spatial and temporal scales. However, our findings show that it is the interplay between the environment and tree form and function that contributes to, and ultimately determines, the geographic location of the species range. At the southern edge of its range, white spruce exhibits unique respiratory responses in the upper vs. lower canopy needles. At the northern limits, the unequal distribution of metabolic activity amongst the different canopy positions is not observed and presumably not needed.

Climate change has, and will continue, to increase air temperature across the white spruce range (Tamarin-Brodsky et al., 2020). This may increase rates of respiration and exacerbate increases in respiratory carbon loss at both the southern and northern range limits of this species. The short-term temperature response curves presented here can be cautiously used to infer how respiration might change with continued warming. We note that thermal acclimation of respiration could directly alter the predicted response (Atkin et al., 2008; Slot & Kitajima, 2015; Vanderwel et al., 2015), most likely lessening the effect of climate warming. However, the noted lack of thermal acclimation of respiration in this species implies that the short-term respiration temperature response may be used to predict the long-term effect of climate warming in white spruce (Benomar et al., 2016). Using the current growing season temperature and the predicted rate of temperature increase by the end of the century (U.S. Climate Resilience Toolkit Climate Explorer), we calculate that white spruce respiration may increase by 67% at the northern range limit regardless of canopy location and by 53% at the southern range limit, where the upper canopy needles are likely to increase more than the lower canopy (46 vs. 60% respectively).

These are massive estimated changes in the leaf carbon flux to the atmosphere that, if unmatched by similar increases in photosynthesis or mediated by thermal acclimation, would have dramatic effects on leaf carbon balance (Ow et al., 2008a; Ow et al., 2008b). Combining the *R*25 rates with these projected rates of increase suggests that any increase in respiration is likely to be more detrimental to the carbon balance of white spruce at the northern relative to the southern range limit. The ecological effects of these physiological changes on the competitiveness of this species and the future species range are unknown but clearly the metabolic adjustments needed to respond to the changing environmental conditions are likely to play a role. We suggest that these effects are at least as severe, if not significantly more so, at the northern end of the range limit than they are at the southern edge. This calls into question the suggestions (ACIA 2005; Zhang et al. 2013; Pearson et al. 2013) that species migrations will shift species ranges northward, and more specifically, to move arctic treeline north in response to climate change.

## Conclusions

The natural distribution of white spruce extends not just across strong environmental gradients, but also to the very limit of tree growth form. North of its current range limit, not only is white spruce not found, but neither are trees of any species. The results presented here may help explain this observation. We clearly show that respiration rates of white spruce at their northern range limit are not only higher than those from trees growing at the southern range limit, but that these rates are extreme. Furthermore, we show that canopy position has a strong influence on the distribution of respiratory activity at the southern but not at the northern range limit, a pattern that likely reflects differences in metabolic activity driven by light. The model of Heskel et al. (2016) was successfully fit to our data and demonstrates that the temperature response of respiration at the southern end of the species distribution is statistically indistinguishable from the NLEv tree plant functional type used in their global survey. In contrast, the northern range limit trees have much higher rates of respiration and differ significantly from the NLEv PFT. Our work supports the conclusion of Griffin et al. (2021), that white spruce needles at the northern edge of the species range respire at what are likely near the species limits, and that this carbon cost likely contributes to the location of northern treeline. Using the short-term temperature response curves to constrain the potential response of respiration to predicted end-of-the-century warming, we show that respiration will have a significant impact on leaf carbon balance that is likely to contribute to future range limits of this species. Our findings raise questions regarding the assertion that species like white spruce will simply shift their range distributions northward in response to warming and suggest that, without a detailed understanding of the myriad ways photosynthesis, respiration and growth will respond to changing climatic conditions, range shifts will be difficult to predict.

## Author Contributions

K.L.G., S.C.S., N.B., L.A.V. and J.U.H.E. designed the research. They were assisted in data collection by Z.M.G. & S.G.B., S.C.S, Z.M.G. & K.L.G. analyzed the data. K.L.G wrote the first draft of the manuscript. All authors contributed to the revisions, editing and submission of the final manuscript.

## Acknowledgements

We thank: Matt Brady and the Black Rock Forest staff, Sarah Sackett and the NASA ABoVE support team, and the Toolik Lake Field Station Staff, for logical and field support. We thank Professor Duncan Menge for advice regarding the interpretation of the nitrogen isotope data and Professor Mary Heskel for helpful discussions about modeling the respiratory temperature response. We thank the two anonymous reviewers and the editor of PCE for their valuable comments and feedback which lead to the significant revision and re-interpretation of this work. This work was supported by NASA ABoVE grant NNX15AT86A and the Arctic LTER (NSF Grant No. 1637459).

## Supplementary information

**Table S1.** Data used to summarize of the relationship between white spruce (<u>Picea glauca</u>) R25 (leaf respiration measured at 25 °C), and the average growing season temperature of the study locations. Entries from the GlobResp database are cited as Atkin et al., 2015 and the original data source can be found there. Where respiration was reported at a temperature other than 25 °C, the given Q10 relationships were used to determine R25. Growing season temperature for the three study sites of McNown & Sullivan (2013), were taken from Sullivan et al. (2021).

| Site | Latitude<br>(°N) | Longitude<br>(°W) | JJA temp<br>(°C) | $R_{25}$<br>($\mu\text{mol m}^{-2} \text{s}^{-1}$) | citation |
| --- | --- | --- | --- | --- | --- |
| BRF, NY | 41.40 | 74.07 | 21.2 | 0.75 | *this study* |
| Ste-Rose de Watford | 46.30 | 70.40 | 15.6 | 2.10 | Benomar et al. 2015 |
| Squatec | 47.84 | 68.52 | 13.9 | 2.60 | Benomar et al. 2015 |
| DeRobidouxville | 48.62 | 71.73 | 13.0 | 2.40 | Benomar et al. 2015 |
| Lac Bergeron | 49.65 | 72.41 | 13.8 | 3.31 | Benomar et al. 2018 |
| Rousseau | 49.15 | 79.20 | 14.9 | 3.37 | Benomar et al. 2018 |
| Deville | 48.62 | 65.72 | 12.4 | 3.31 | Benomar et al. 2018 |
| Asselin | 47.84 | 68.52 | 14.0 | 2.88 | Benomar et al. 2018 |
| Picard | 47.34 | 73.57 | 14.4 | 3.72 | Benomar et al. 2018 |
| Dorion | 46.30 | 70.40 | 13.9 | 2.32 | Benomar et al. 2018 |
| Watford | 46.05 | 76.26 | 16.6 | 2.97 | Benomar et al. 2018 |
| Wendover | 45.98 | 72.52 | 15.9 | 2.34 | Benomar et al. 2018 |
| Terrace, AK | 67.47 | 162.22 | 14.3 | 2.28 | McNown & Sullivan 2013 |
| Forest, AK | 67.47 | 162.22 | 14.3 | 2.25 | McNown & Sullivan 2013 |
| Treeline, AK | 67.47 | 162.22 | 14.3 | 3.62 | McNown & Sullivan 2013 |
| Reichetal Wisconsin | 42.50 | 90.00 | 20.7 | 1.14 | Atkin 2015 |
| Minnesota | 46.17 | 92.87 | 17.7 | 0.93 | Lusk & Reich |
| CloquetFC | 46.71 | 92.53 | 17.7 | 0.81 | Reich et al 2016 |
| B4W-HWRC-OC | 47.94 | 96.33 | 17.3 | 0.75 | Atkin 2015 |
| FTE, AK | 67.98 | 149.75 | 14.8 | 2.35 | *this study* |

Note while several other publications report respiration rates for *Picea glauca* leaves, they do not provide all of the information needed for this analysis.

**Figure S1.**
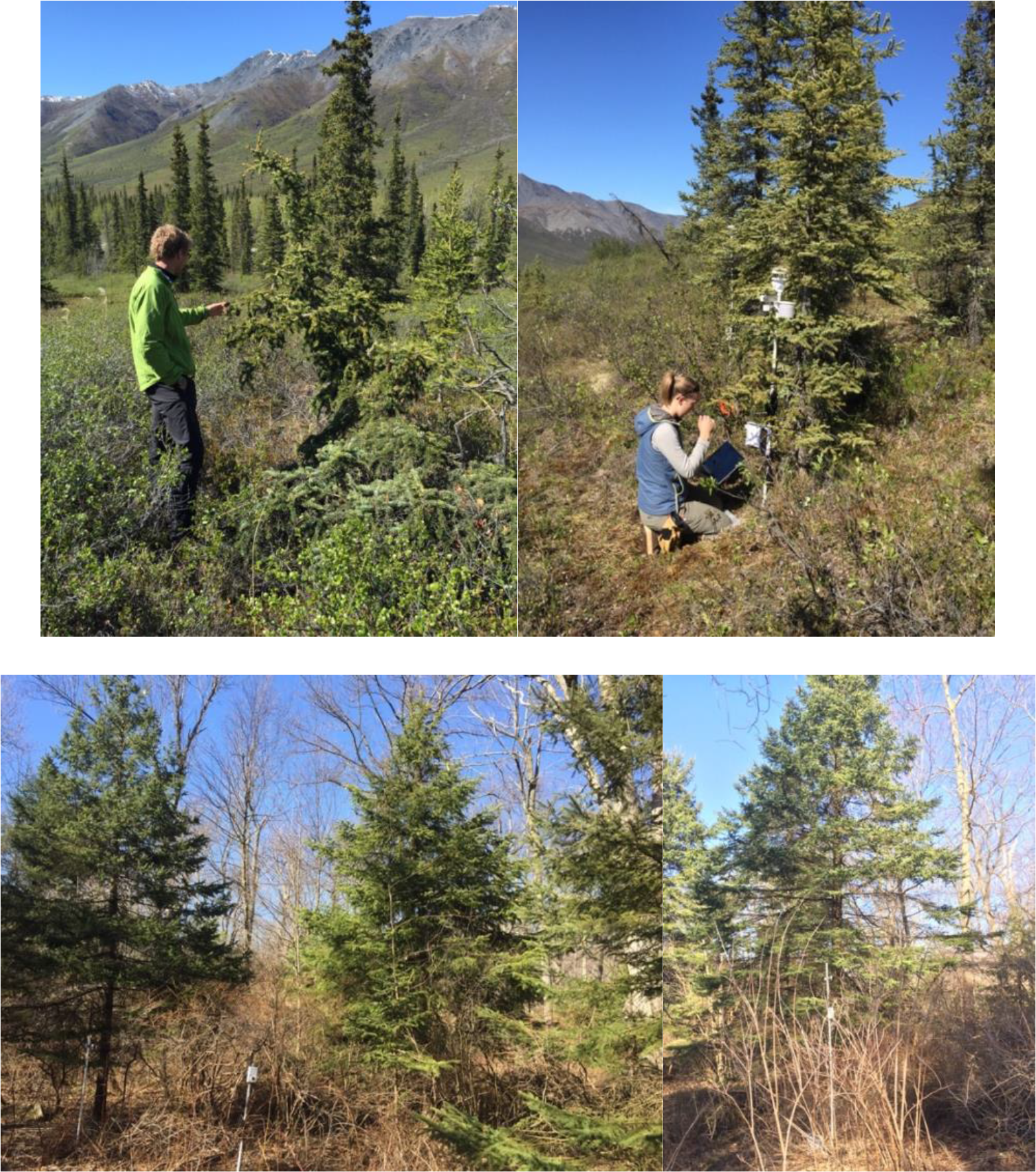
Photos of the tree canopies showing the open nature of the northern treeline site (top) vs. the more closed canopy of the southern site (bottom). Note the southern site photo was taken in Spring, and thus the surrounding deciduous canopy is not present.

**Figure S2.**
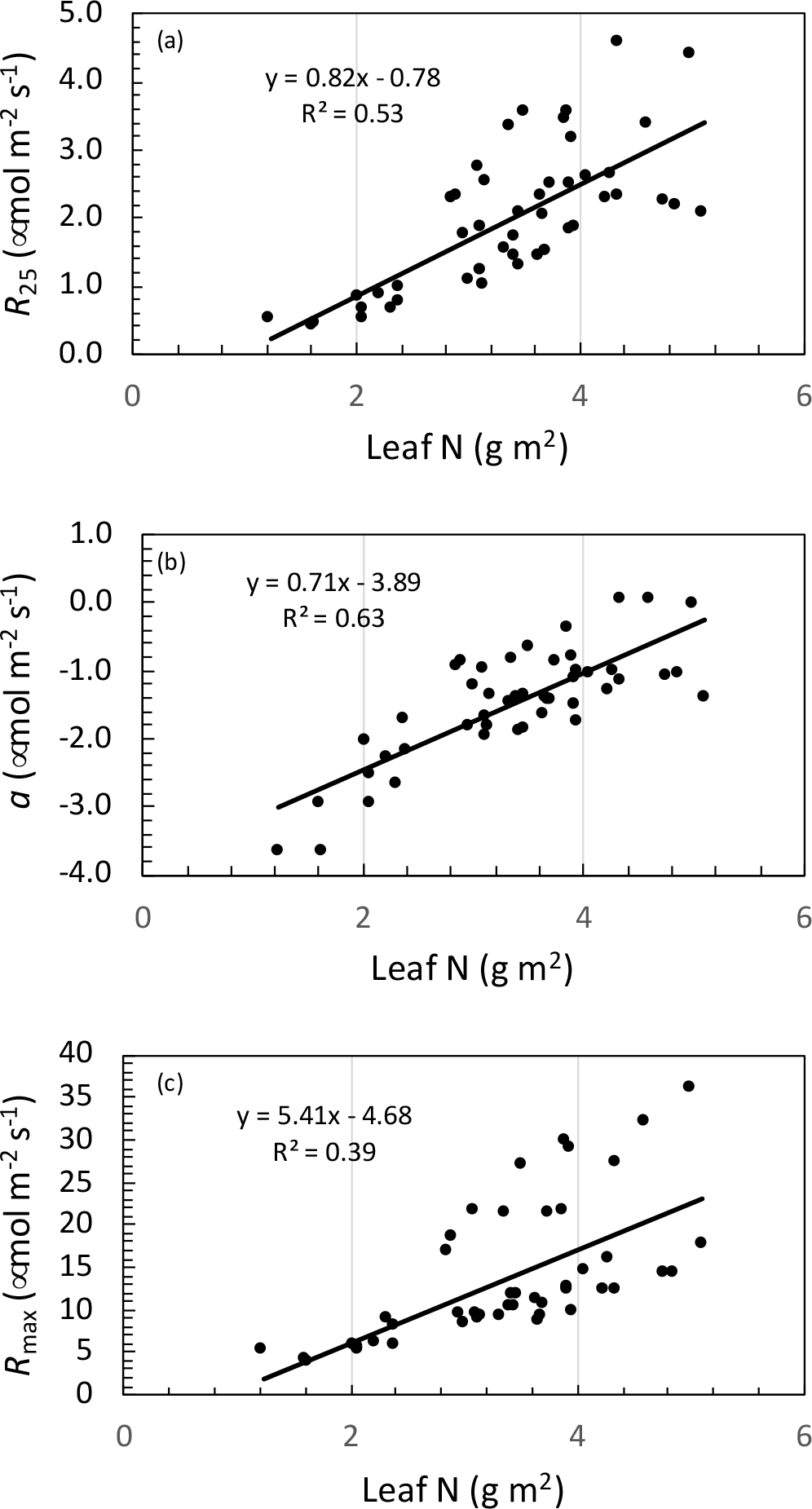
Relationships between leaf nitrogen and respiration of white spruce (*<u>Picea glauca</u>*) foliage. Panel (a) is the relationship between respiration measured at a leaf temperature of 25 °C and expressed on a projected leaf area basis and leaf nitrogen per m^-2^. Panel (b) is the *a* coefficient of the polynomial model (Heskel et al., 2016) fit to the measured high-resolution leaf respiration temperature response curves vs. leaf nitrogen per m^-2^. Panel (c) is the maximum rate of respiration from a temperature response curve, expressed on a leaf area basis. All regressions are statistical significance (p<0.05), n = 24.

